# Omicron variant of SARS-CoV-2 exhibits an increased resilience to the antiviral type I interferon response

**DOI:** 10.1101/2022.01.20.476754

**Authors:** Lyudmila Shalamova, Ulrike Felgenhauer, Andreas R. Schaubmar, Kathrin Büttner, Marek Widera, Sandra Ciesek, Friedemann Weber

**Affiliations:** Institute for Virology, FB10-Veterinary Medicine, Justus-Liebig University; D-35392 Giessen, Germany; Unit for Biomathematics and Data processing, FB10-Veterinary Medicine, Justus-Liebig University; D35392 Giessen, Germany; Institute for Medical Virology, University Hospital, Goethe University; Frankfurt am Main, Germany; German Centre for Infection Research (DZIF), partner sites Giessen and Frankfurt, Germany; Branch Translational Medicine and Pharmacology, Fraunhofer Institute for Molecular Biology and Applied Ecology (IME), 60596 Frankfurt am Main, Germany

**Keywords:** COVID-19, SARS-CoV-2, Omicron variant, interferon induction, interferon sensitivity

## Abstract

The new variant of concern (VOC) of the severe acute respiratory syndrome coronavirus 2 (SARS-CoV-2), Omicron (B.1.1.529), is genetically very different from other VOCs. We compared Omicron with the preceding VOC Delta (B.1.617.2) and the wildtype strain (B.1) with respect to their interactions with the antiviral type I interferon (IFN-alpha/beta) response in infected cells. Our data indicate that Omicron has gained an elevated capability to suppress IFN-beta induction upon infection and to better withstand the antiviral state imposed by exogenously added IFN-alpha.

## Introduction

Omicron (B.1.1.529) is a new variant of concern (VOC) of the pandemic severe acute respiratory syndrome coronavirus 2 (SARS-CoV-2) causing COVID-19. Since its detection in South Africa in November 2021 (1), Omicron has rapidly spread in all countries where it was introduced, indicating elevated infectivity and a certain resistance to pre-existing immunity (2). These features are due to an unprecedented number of mutations that are distinguishing Omicron from the original coronavirus that emerged in Wuhan, China, at the end of 2019, as well as from the subsequently appearing VOCs like e.g. Delta (B.1.617.2) (1). Omicron exhibits a high resistance to neutralization by antibodies directed against the previously dominant lineages (3), an increased binding to the main virus receptor ACE2 (4), and a switch to the endosomal entry route (5). Moreover, it replicated more efficiently in cells of the upper respiratory tract (5), most likely contributing to its hyper-transmissibility (2).

Type I interferons (IFN-alpha/beta) constitute the first line of the innate immune defense against invading viruses (6). Upon infection, viral hallmark structures like e.g. doublestranded RNA are recognized by cellular sensors that, in turn, initiate a so-called antiviral signaling chain culminating in the induction of genes for IFN-beta and other cytokines. Secreted IFN then binds to its cognate receptor in an autocrine and paracrine fashion to establish an antiviral state in the cell. SARS-CoV-2, like other highly pathogenic viruses (6), therefore evolved a series of countermeasures that suppress induction of IFNs or the IFN-stimulated signaling (7). Despite the more than 15 viral proteins exhibiting IFN-antagonistic activity, infection by SARS-CoV-2 still induces a certain level of type I IFNs and other cytokines (8) and exogenously added IFN is inhibitory to viral replication (8–10).

## Results

We investigated the capability of Omicron to interact with the antiviral type I IFN system, in comparison to the parental “wildtype” (wt) SARS-CoV-2 strain (B.1) and the still most prevalent VOC, Delta (B.1.617.2). As a first step, induction of innate immunity was measured in the Calu-3 human lung epithelial cell model using RT-qPCR. As representative marker genes, we chose immediate-early virus response cytokines IFN-beta, IFN-lambda1, and CXCL10 (11), and as positive control for innate immunity marker gene activation we used the IFN-inducing Rift Valley fever virus mutant Clone 13 (11). Figure 1A shows that all SARS-CoV-2 strains are inducing the cytokine genes, but as expected their induction levels were much lower than for the control virus Clone 13. Strikingly, however, Omicron tended to exhibit the lowest activation levels regarding the three cytokines, especially IFN-beta. Of note, levels of viral RNAs were comparable for all SARS-CoV-2 strains, excluding the possibility that Omicron simply generated less IFN-inducing RNA than the other coronaviruses (Fig. 1B).

**Figure 1.**
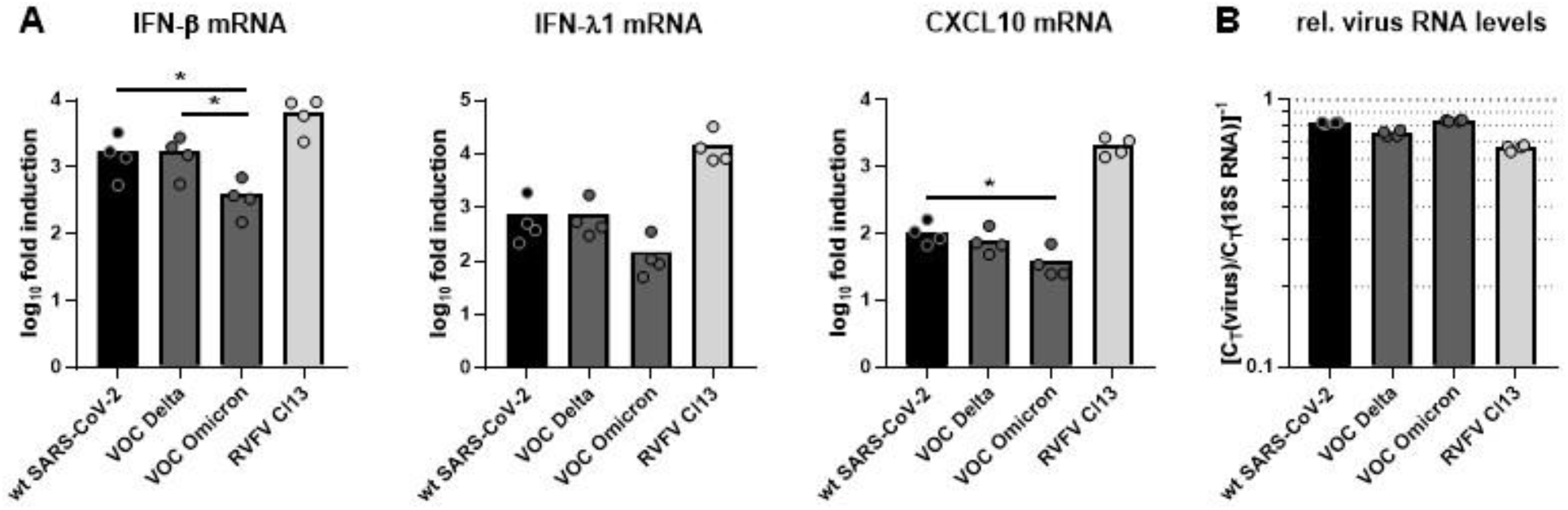
Cytokine induction by strains of SARS-CoV-2. Human Calu-3 cells were infected with the indicated viruses at a multiplicity of infection (MOI) of 1, lysed at 24 h post-infection, and the extracted RNA was subjected to RT-qPCR analyses for mRNAs of cytokines (**A**). Results are given as fold-induction over the uninfected mock control, normalized to 18S rRNA. (**B**) Viral gene expression depicted as C_T_ values for viral E (coronaviruses) or L (RVFV Cl13) gene normalized to 18S rRNA control. Individual data points (dots) and mean values (bars) are shown. The log_10_ transformed values of the different coronaviruses were pairwise tested by one-factorial ANOVA with Bonferroni correction. Asterisk (*) indicates p < 0.05. All other comparisons had p values > 0.05 (not shown). Note that the internal control for IFN induction, RVFV Cl13, was not included in the statistical testing.

The results of the induction experiments suggest that Omicron has an improved ability to suppress the induction of antiviral cytokines. In order to investigate the other end of the IFN response, namely sensitivity to exogenously added IFNs, we pre-treated cells with increasing amounts of IFN-alpha, infected them with the various SARS-CoV-2 strains at a low MOI, and measured virus yields 24 h later. Figure 2 (A: absolute titers, B: relative titers) shows that all viruses were reduced by IFN in a dose-dependent manner. However, 50 units IFN/ml were able to suppress titers of wt SARS-CoV-2 and Delta by 10-fold or more, but Omicron titers remained within the same order of magnitude. In fact, 500 units IFN/ml were required to suppress Omicron to a similar extent as 50 IFN/ml did for the wt, and 100 units IFN/ml did for Delta. In order to compare the dose-response slopes of the individual viruses, linear regressions on the log_10_-transformed data were performed (Fig. 3). These slopes were then compared using a covariance analysis with the individual virus as fixed effect and the dose of IFN-alpha as covariable. All pairwise comparisons were adjusted using the Bonferroni correction. The linear regressions already showed that the IFN dose responses by wt SARS-CoV-2 and Delta had steeper slopes than Omicron (see Figs. 2A and 3). Indeed, the pairwise comparisons (Table 1) revealed that there was no significant difference in the slopes between wt SARS-CoV-2 and Delta, whereas the slopes of wt SARS-CoV-2 vs. Omicron as well as of Delta vs. Omicron were significantly different. These data indicate that Omicron has gained an increased resistance to human type I IFNs.

**Figure 2.**
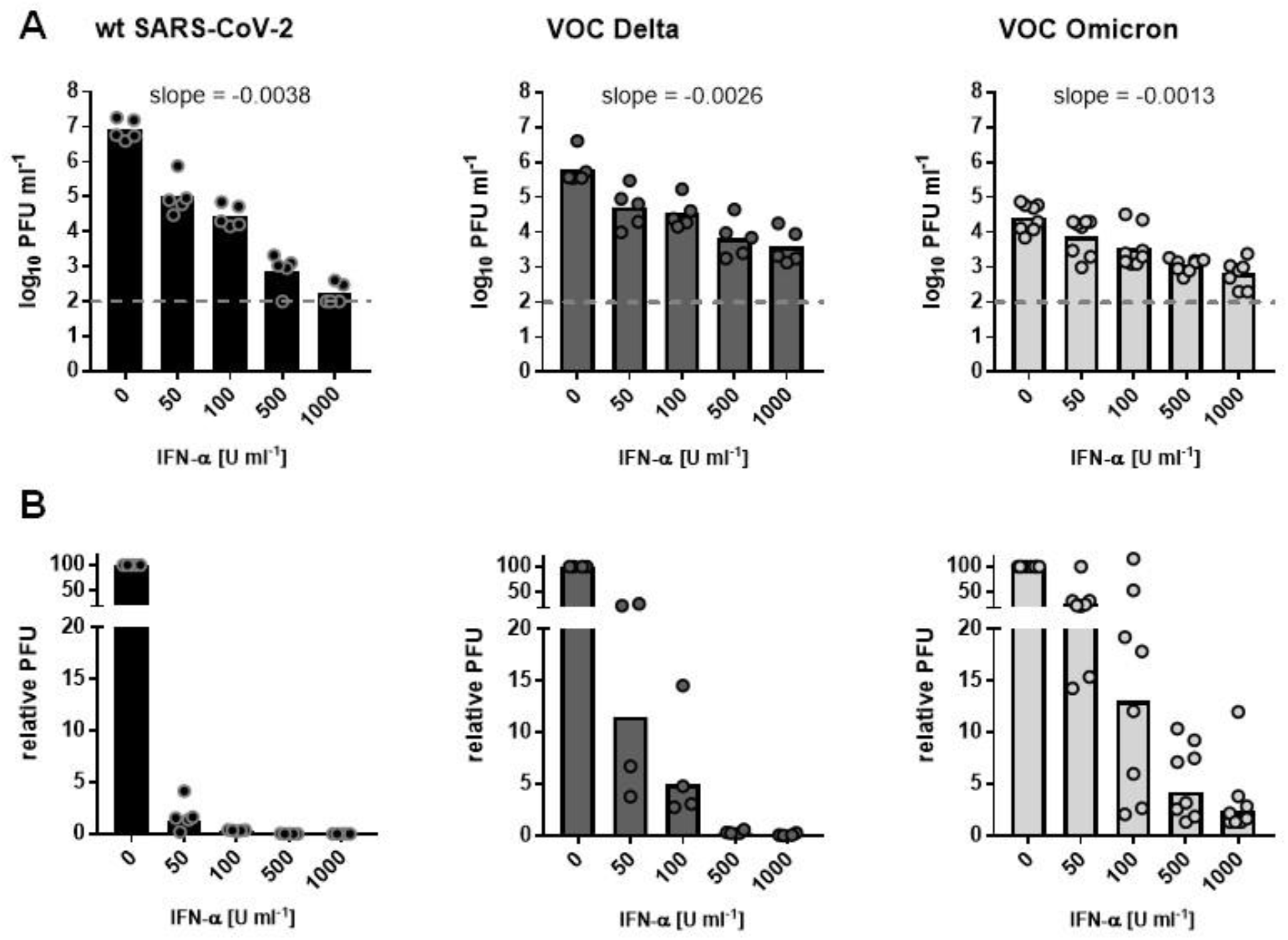
Sensitivity to type I IFN dose escalation. Calu-3 cells were pre-treated with increasing amounts of recombinant human IFN-alpha for 24 h prior to infection with the indicated viruses at an MOI of 0.01 for 24 h. (A) raw virus titer data, (B) data normalized to virus titers of the non-treated control. Titer values that were below the plaque assay detection level (100 PFU/ml; dashed line) were set to 100 PFU/ml. Slopes were calculated by linear regressions on the log_10_-transformed data.

**Figure 3.**
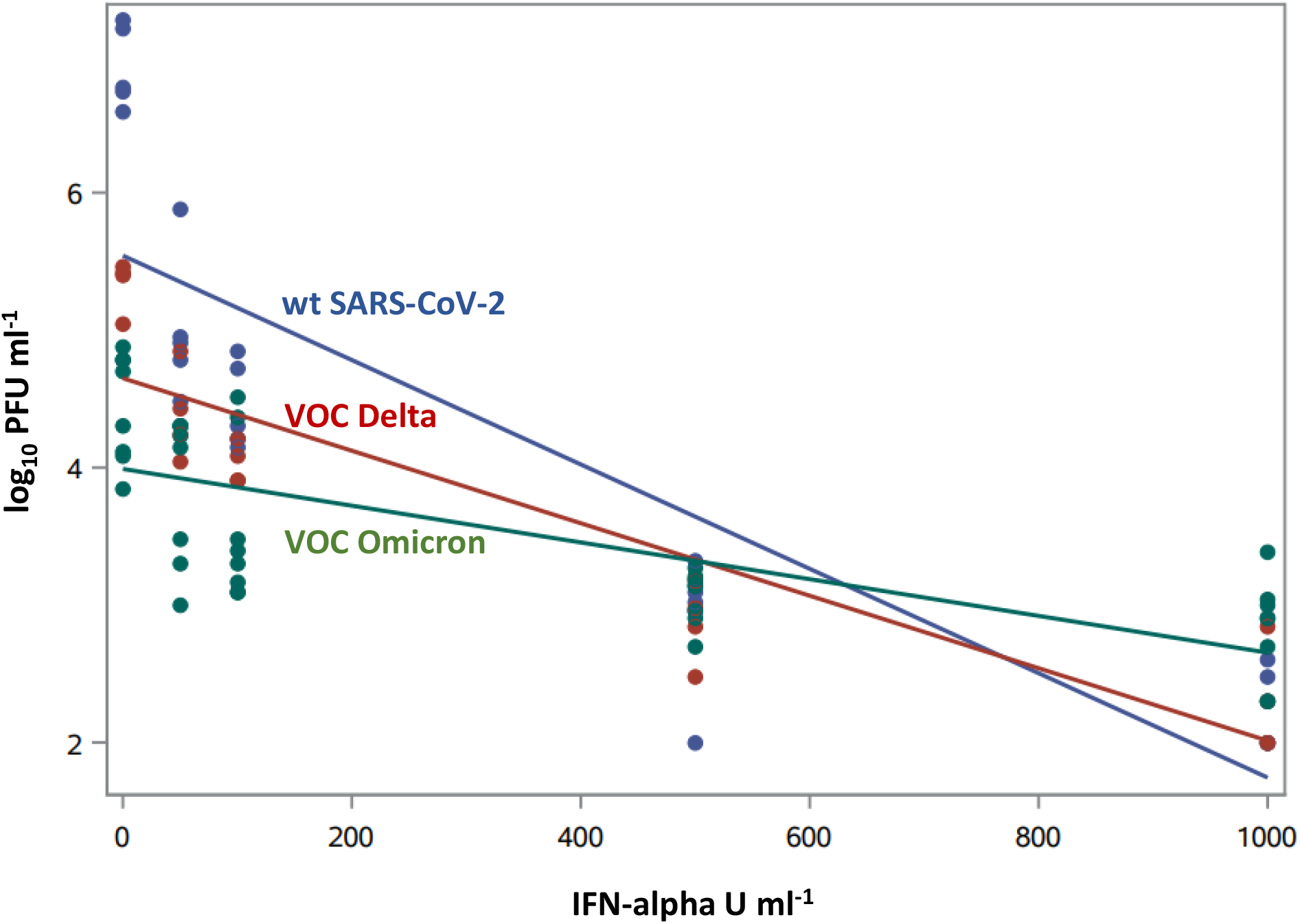
Linear regression of the three SARS-CoV-2 viruses in dependence of the IFN dose. Analysis is based on the log_10_-transformed data shown in Fig. 2A

**Table 1.** Pairwise comparisons of IFN sensitivity regression slopes (covariance analysis with Bonferroni correction).

| **Comparison** | **Bonferroni-adjusted p** |
| --- | --- |
| wt SARS-CoV-2 vs. VOC Delta | 0.0871 |
| wt SARS-CoV-2 vs. VOC Omicron | < 0.0001 (*) |
| VOC Delta vs. VOC Omicron | 0.0234 (*) |

## Discussion

Exogenously added IFNs were shown to restrict the original wt SARS-CoV-2 in cell culture (8–10), and in patients an early type I IFN therapy is associated with reduced mortality (12). This may also apply to the IFN that is endogenously produced by the infected individual, as the strength of the IFN response correlates with disease severity. In children, a pre-activated IFN system that enables a rapid and strong antiviral response is controlling infection (13), whereas in the elderly the IFN system is less active and can be additionally hampered by neutralizing anti-IFN autoantibodies (14). Thus, the ability of a virus to suppress IFN induction and antiviral IFN action are important determinants of COVID-19 risk, and the outcome of infection is dependent on the timing, amount and localization of IFN production. Our results are in line with reports that the earlier VOC Alpha suppresses IFN induction more effectively than preceding wt SARS-CoV-2 isolates (15) and that the more pathogenic SARS-CoV-1 from 2003 is less IFN-sensitive than wt SARS-CoV-2 (9, 10). Thus, IFN escape is an additional feature besides antibody escape, increased ACE2 affinity and endosomal entry that may contribute to the hyper- transmissibility of Omicron.

## Materials and Methods

### Cells and viruses

Calu-3, Caco-2, Vero E6 and Vero E6-TMPRSS2 cells (kindly provided by Stefan Poehlmann, German Primate Center, Goettingen) were cultivated in Dulbecco’s modified Eagle’s medium (DMEM) supplemented with 10% fetal bovine serum. SARS-CoV-2 strains “wildtype (wt)” (SARS-CoV-2/München-1.2/2020/984 (B.1) p.4; (16), (kindly provided by Christian Drosten, Charité, Berlin)), “VOC Delta” (SARS-CoV-2 B.1.617.2 (FFM-IND8424/2021) (17)) and “VOC Omicron” (SARS-CoV-2 B.1.1.529 (FFM- SIM0550/2021) (3)) were grown on Vero E6-TMPRSS2 cells (wt SARS-CoV-2) or Caco- 2 cells (VOCs Delta, Omicron), respectively and titrated on Vero E6 cells. Infection experiments were done under biosafety level 3 conditions. Of note, all cells and viruses were tested mycoplasma-negative.

### Interferon induction analysis

Calu-3 cells were seeded into 24-well plates (7.5×10E4 per well) and infected 24 h later at an MOI of 1. At 24 h post-infection, cells were lysed in RLT buffer and total RNA was extracted using the RNeasy mini kit (Qiagen). RNA was reverse transcribed with the prime Script RT reagent kit (Takara). Differential regulation of cellular genes was assessed using TB Green™ Premix Ex Taq™ II (Tli RNase H Plus; Takara) according to manufacturer’s instructions with QuantiTect primer assays (Qiagen; Hs_IFNB1_1_SG, QT00203763; Hs_IFNL1_2_SG, QT01033564; Hs_CXCL10_1_SG, QT01003065). Viral RNA loads in samples were measured using Premix Ex Taq (probe qPCR; Takara) with primers and probes as follows: for SARS-CoV E gene, E_Sarbeco_F: 5’-ACAGGTACGTTAATAGTTAATAGCGT-3’, E_Sarbeco_P1: 5’-FAM-ACACTAGCCATCCTTACTGCGCTTCG-BBQ- 3’, E_Sarbeco_R: 5’-ATATTGCAGCAGTACGCACACA-3’ (18); for RVFV L segment, fwd, 5’- TGAAAATTCCTGAGACACATGG-3’, rev, 5’-ACTTCCTTGCATCATCTGATG-3’, probe, 5’-6-FAM-CAATGTAAGGGGCCTGTGTGGACTTGTG-BHQ1-3’ (19). Fold induction was calculated according to the threshold cycle (ΔΔCT) method using 18S rRNA as a reference gene.

### Interferon treatment

Cells seeded into 24-well plates (7.5×10E4 per well) were pre-treated for 24 h with 50, 100, 500 or 1000 U/ml pan-species IFN-alpha (B/D) (PBL Assay Science) and infected at an MOI of 0.01. At 24 h post-infection, cell supernatants were collected and titrated by plaque assay on Vero E6 cells.

### Statistical analyses

Statistical analyses were done using the program package SAS 9.4.

## Acknowledgments

We thank Christian Drosten and Stefan Poehlmann for kindly providing wt SARS-CoV-2 and Vero E6-TMPRSS2 cells, respectively. Work in the authors’ laboratories was funded by the Deutsche Forschungsgemeinschaft (DFG) Clinical Research Unit KFO309 Project P1, 284237345 (FW), the Bundesministerium für Bildung und Forschung (BMBF) RAPID grant 01KI1723E (FW), the European Union’s Horizon 2020 research and innovation program under grant agreement no. 101005026 (MAD-CoV-2) (FW), and the Pandemie Netzwerk of the Land Hesse (SC, FW)

## References

1. R. Viana et al., Rapid epidemic expansion of the SARS-CoV-2 Omicron variant in southern Africa. medRxiv 10.1101/2021.12.19.21268028, 2021.2012.2019.21268028 (2021).

2. F. Grabowski, M. Kochańczyk, T. Lipniacki, Omicron strain spreads with the doubling time of 3.2—3.6 days in South Africa province of Gauteng that achieved herd immunity to Delta variant. medRxiv 10.1101/2021.12.08.21267494, 2021.2012.2008.21267494 (2021).

3. A. Wilhelm et al., Reduced Neutralization of SARS-CoV-2 Omicron Variant by Vaccine Sera and monoclonal antibodies. medRxiv 10.1101/2021.12.07.21267432, 2021.2012.2007.21267432 (2021).

4. M. McCallum et al., Structural basis of SARS-CoV-2 Omicron immune evasion and receptor engagement. bioRxiv 10.1101/2021.12.28.474380, 2021.2012.2028.474380 (2021).

5. T. P. Peacock et al., The SARS-CoV-2 variant, Omicron, shows rapid replication in human primary nasal epithelial cultures and efficiently uses the endosomal route of entry. preprint (2022).

6. A. Garcia-Sastre, Ten Strategies of Interferon Evasion by Viruses. Cell Host Microbe 22, 176–184 (2017).

7. E. Palermo, D. Di Carlo, M. Sgarbanti, J. Hiscott, Type I Interferons in COVID-19 Pathogenesis. Biology-Basel 10(2021).

8. A. Banerjee et al., Experimental and natural evidence of SARS-CoV-2-infection-induced activation of type I interferon responses. iScience 24, 102477 (2021).

9. U. Felgenhauer et al., Inhibition of SARS-CoV-2 by type I and type III interferons. J Biol Chem 295, 13958–13964 (2020).

10. K. G. Lokugamage et al., Type I Interferon Susceptibility Distinguishes SARS-CoV-2 from SARS-CoV. J Virol 94 (2020).

11. M. Holzer et al., Virus- and Interferon Alpha-Induced Transcriptomes of Cells from the Microbat Myotis daubentonii. iScience 19, 647–661 (2019).

12. N. Wang et al., Retrospective Multicenter Cohort Study Shows Early Interferon Therapy Is Associated with Favorable Clinical Responses in COVID-19 Patients. Cell Host Microbe 28, 455–+ (2020).

13. J. Loske et al., Pre-activated antiviral innate immunity in the upper airways controls early SARS-CoV-2 infection in children. Nat Biotechnol 10.1038/s41587-021-01037-9 (2021).

14. P. Bastard et al., Autoantibodies neutralizing type I IFNs are present in ~4% of uninfected individuals over 70 years old and account for ~20% of COVID-19 deaths. Sci Immunol 6(2021).

15. L. G. Thorne et al., Evolution of enhanced innate immune evasion by the SARS-CoV-2 B.1.1.7 UK variant. bioRxiv 10.1101/2021.06.06.446826, 2021.2006.2006.446826 (2021).

16. C. Rothe et al., Transmission of 2019-nCoV Infection from an Asymptomatic Contact in Germany. New Engl J Med 382, 970–971 (2020).

17. A. Wilhelm et al., Antibody-Mediated Neutralization of Authentic SARS-CoV-2 B.1.617 Variants Harboring L452R and T478K/E484Q. Viruses 13(2021).

18. V. M. Corman et al., Detection of 2019 novel coronavirus (2019-nCoV) by real-time RT-PCR. Euro Surveill 25 (2020).

19. B. H. Bird, D. A. Bawiec, T. G. Ksiazek, T. R. Shoemaker, S. T. Nichol, Highly sensitive and broadly reactive quantitative reverse transcription-PCR assay for high-throughput detection of Rift Valley fever virus. J Clin Microbiol 45, 3506–3513 (2007).

